# Spatial and Temporal Organization of Composite Receptive Fields in the Songbird Auditory Forebrain

**DOI:** 10.1101/2021.07.21.453115

**Authors:** Nasim W. Vahidi

**Affiliations:** Department of Electrical and Computer Engineering, University of California San Diego, San

**Keywords:** composite receptive fields, spatial and temporal map, neural decoding, stimuli encoding, auditory

## Abstract

The mechanisms underlying how single auditory neurons and neuron populations encode natural and acoustically complex vocal signals, such as human speech or bird songs, are not well understood. Classical models focus on individual neurons, whose spike rates vary systematically as a function of change in a small number of simple acoustic dimensions. However, neurons in the caudal medial nidopallium (NCM), an auditory forebrain region in songbirds that is analogous to the secondary auditory cortex in mammals, have composite receptive fields (CRFs) that comprise multiple acoustic features tied to both increases and decreases in firing rates. Here, we investigated the anatomical organization and temporal activation patterns of auditory CRFs in European starlings exposed to natural vocal communication signals (songs). We recorded extracellular electrophysiological responses to various bird songs at auditory NCM sites, including both single and multiple neurons, and we then applied a quadratic model to extract large sets of CRF features that were tied to excitatory and suppressive responses at each measurement site. We found that the superset of CRF features yielded spatially and temporally distributed, generalizable representations of a conspecific song. Individual sites responded to acoustically diverse features, as there was no discernable organization of features across anatomically ordered sites. The CRF features at each site yielded broad, temporally distributed responses that spanned the entire duration of many starling songs, which can last for 50 s or more. Based on these results, we estimated that a nearly complete representation of any conspecific song, regardless of length, can be obtained by evaluating populations as small as 100 neurons. We conclude that natural acoustic communication signals drive a distributed yet highly redundant representation across the songbird auditory forebrain, in which adjacent neurons contribute to the encoding of multiple diverse and time-varying spectro-temporal features.

## 1 Introduction

The sensory encoding mechanisms that underlie the neural representation of complex, natural, acoustic communication signals, such as human speech or bird songs, are poorly understood. The classical model for single sensory neuron responses, the receptive field, has had a long and productive history (Hartline, 1938; Hubel and Wiesel, 1959; Quiroga et al., 2005). In the sensory space, receptive fields of neurons are responsible for this encoding mechanism (Wu et al., 2002). In a previous study, we showed that single neurons in secondary auditory cortex-like regions of European starlings (*Sturnus vulgaris*) have composite receptive fields (CRFs) (Kozlov and Gentner, 2016). Unlike linear receptive fields (LRFs), CRFs contain multiple, distinct receptive field features.

In this study, we created a spatiotemporal map of CRFs by generating a pool of CRFs from a large population of cells in the higher auditory region of starling brains. We then used this map to investigate auditory processing, natural song encoding, and neural response decoding mechanisms in birdsong. “Stimuli encoding” refers to the process of mapping stimuli to responses and is primarily used to reconstruct stimuli from neural responses (Mesgarani et al., 2009; Pasley et al., 2012). In contrast, “neural decoding” is a reverse mapping process that maps responses to stimuli and is used to reconstruct responses from stimuli (Koyama, 2012). Neural decoding is mainly used to predict brain responses and has applications for brain-computer interfaces (Velliste et al., 2008; Yildiz et al., 2016).

Existing models for creating receptive fields of neurons include the spike-triggered average (STA), which builds spectro-temporal receptive fields (de Boer and Kuyper, 1968), as well as the spike-triggered covariance (STC) (Schwartz et al., 2002) and the maximally informative dimensions (MID) (Sharpee et al., 2004). Although these models have provided a wealth of information about receptive fields and the characteristics of stimuli and responses, they suffer from several drawbacks, including the inability to characterize nonlinear stimulus-response properties (Theunissen et al., 2000), the requirement to work only with natural stimuli (e.g., human speech or bird song) (Eggermont et al., 1983; Schwartz et al., 2006), and the fact that they can only identify a small number of relevant receptive field features with respect to natural stimuli (Kozlov and Gentner, 2016). In this study, we used the maximum noise entropy (MNE) model to overcome most of these limitations (Sharpee et al., 2004; Bialek and de Ruyter van Steveninck, 2005; Fitzgerald et al., 2011b, 2011a; Kozlov and Gentner, 2016; Vahidi et al., 2020). The MNE model looks for mutual information and the highest correlation between auditory stimuli and neural responses in the form of CRFs at the cellular level.

To map CRFs of a population of cells with respect to various stimuli, we used high-density electrodes to record action potentials from the higher-level auditory cortex (the caudomedial nidopallium [NCM]) of five European starlings during playback of starling bird songs (Figure 1A). Our findings might also be applicable to mammals and humans, as birds and mammals process auditory data in a similar way at the cellular level (Karten, 2013; Harris, 2015).

**Figure 1:**
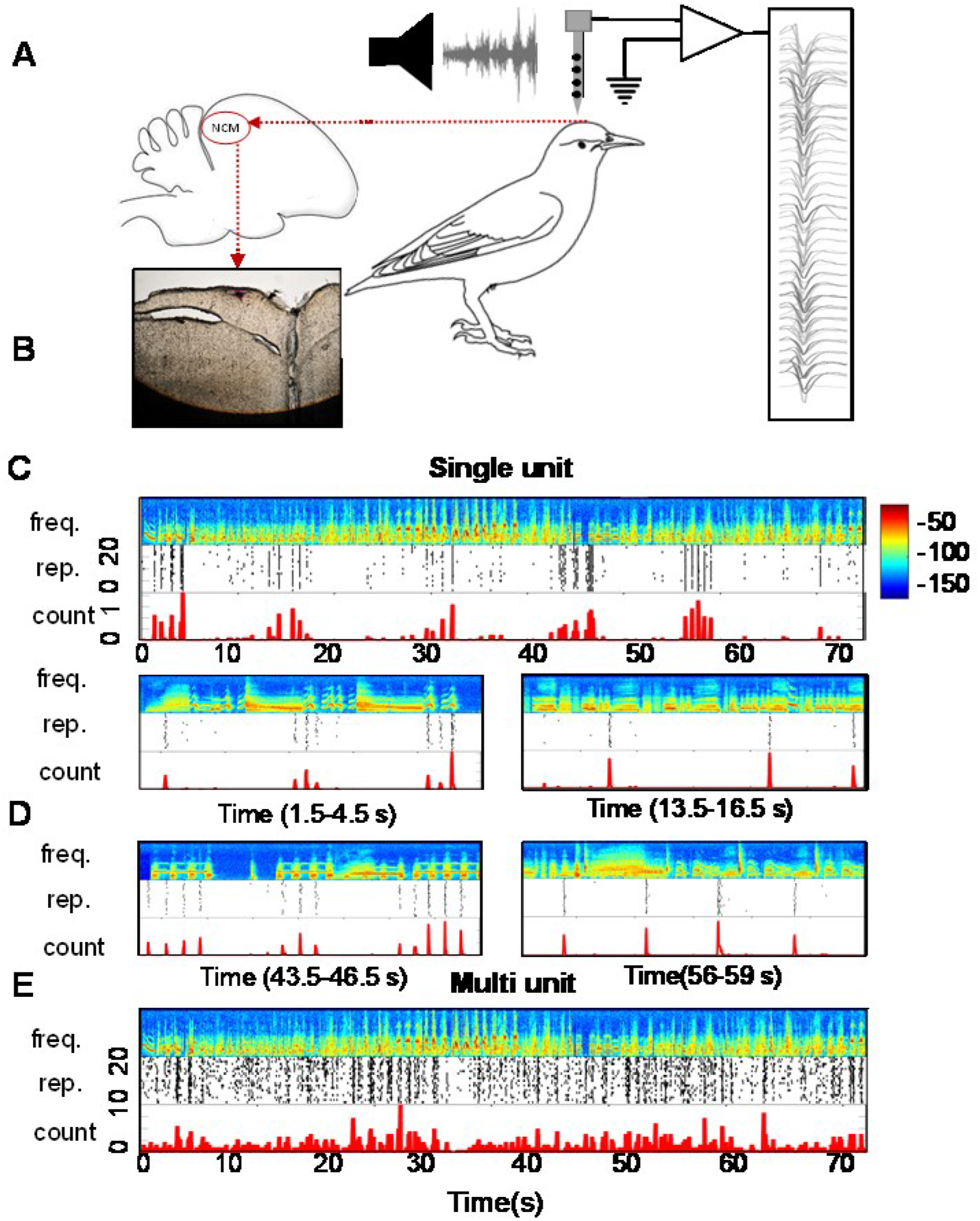
Electrophysiological recording. (A) We implanted high-density silicon recording electrodes in the caudomedial nidopallium (NCM) of anesthetized starlings, then presented conspecific songs while recording evoked extracellular responses (spikes) single and multi-unit neuron clusters. The red arrow demonstrates the location of the implanted electrode. Spike waveforms recorded from cells from subject 4-Pen1 are shown in the rectangular box. (B) Histology of a starling brain (dorsal view). The Red Dil stain shows the location of the implanted array in the NCM. (C) Example response of a well-isolated single neuron, showing a spectrogram for a 70 s of the full stimulus with frequency range of 20 Hz-20 kHz (top; color bar on the right indicates power density), raster plot of stimulus evoked spiking responses across 20 repetitions of the shown stimulus (middle), and the corresponding average spike counts (ca. 20 ms bins) across repetitions. (D) Four separate three-second windows measured from a single cell, showing how various acoustic features are affecting the firing rates of a cell. (E) Example of measurements of multi-unit activity. Figure descriptions for (D) and (E) are the same as for (C).

After sorting and extracting neurons from starling brains, we performed MNE estimation on a population of 154 cells and then extracted the ten most significant facilitatory and ten most significant suppressive CRFs for each cell. Significant CRFs were identified as spikes with either the highest probability of occurrence (facilitatory) or the lowest probability of occurrence (suppressive),as determined by the MNE logistic quadratic model. The first ten significant facilitatory CRFs and the first ten significant suppressive CRFs contained the least amount of noise, on average, across our

154 cells (Figure 2). Using those 20 CRFs per cell, we generated a pool of 3,080 CRFs (20 CRFs * 154 cells) for this study.

**Figure 2:**
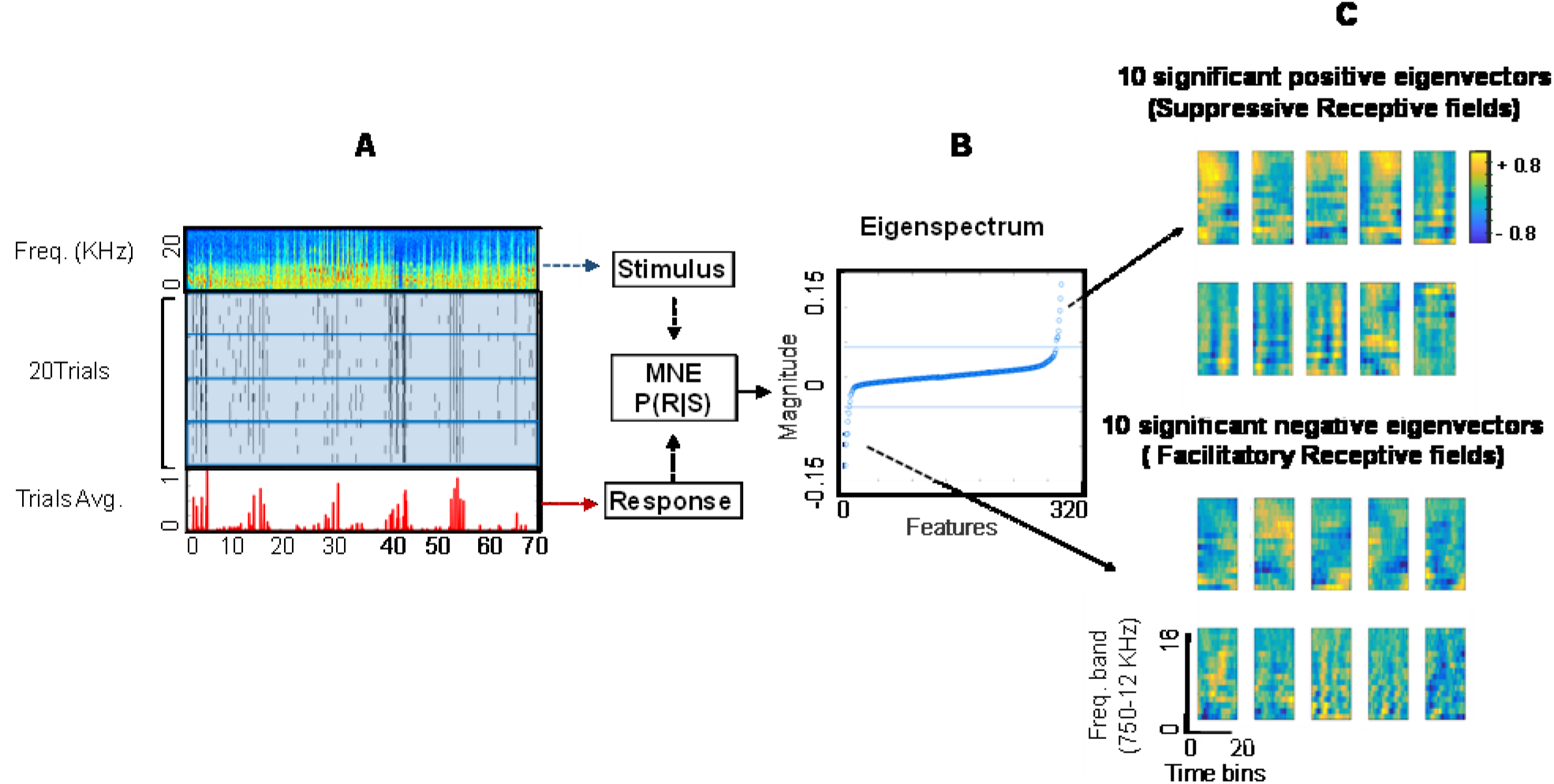
Extracting composite receptive fields (CRFs) from caudomedial nidopallium neurons using the maximum noise entropy (MNE) model. (A) First row: Bird song spectrogram. Middle rows: Raster plot of spiking activities of a single cell measured across 20 trials. Bottom row: Average results across the trials (highlighted in red). (B) Power spectral density of the bird song spectrogram was used as the stimulus input and trials average were used as the response input in the MNE model. The blue curve demonstrates the eigenspectrum (the magnitude of the eigenvalues) of the J-matrix (see section 2.5) for the neuron shown in (A). The dashed lines indicate the largest positive and negative eigenvalues obtained from 500 symmetrical Gaussian random matrices with the same mean and variance as the J-matrix. (C) The eigenvectors corresponding to the 10 most significant positive (top) and negative eigenvalues (bottom) in the J-matrix. We interpret these as the 10 strongest suppressive and 10 strongest facilitatory quadratic features in the neurons’ CRF. Each feature shows the response-relevant power across 16 frequency bands (spanning 750-12 kHz) over 20 time bins (ca. 400 ms total) preceding a change in spike rate. The color bar on the top right shows the power density of the CRF features.

To assess the performance of the MNE model, we tested the model on our population of neurons by predicting the neural spike train response to various novel songs. Our results showed that neural response prediction using CRFs was highly accurate (Figure 4), which indicates that the MNE model is a good candidate for neural decoding.

**Figure 3:**
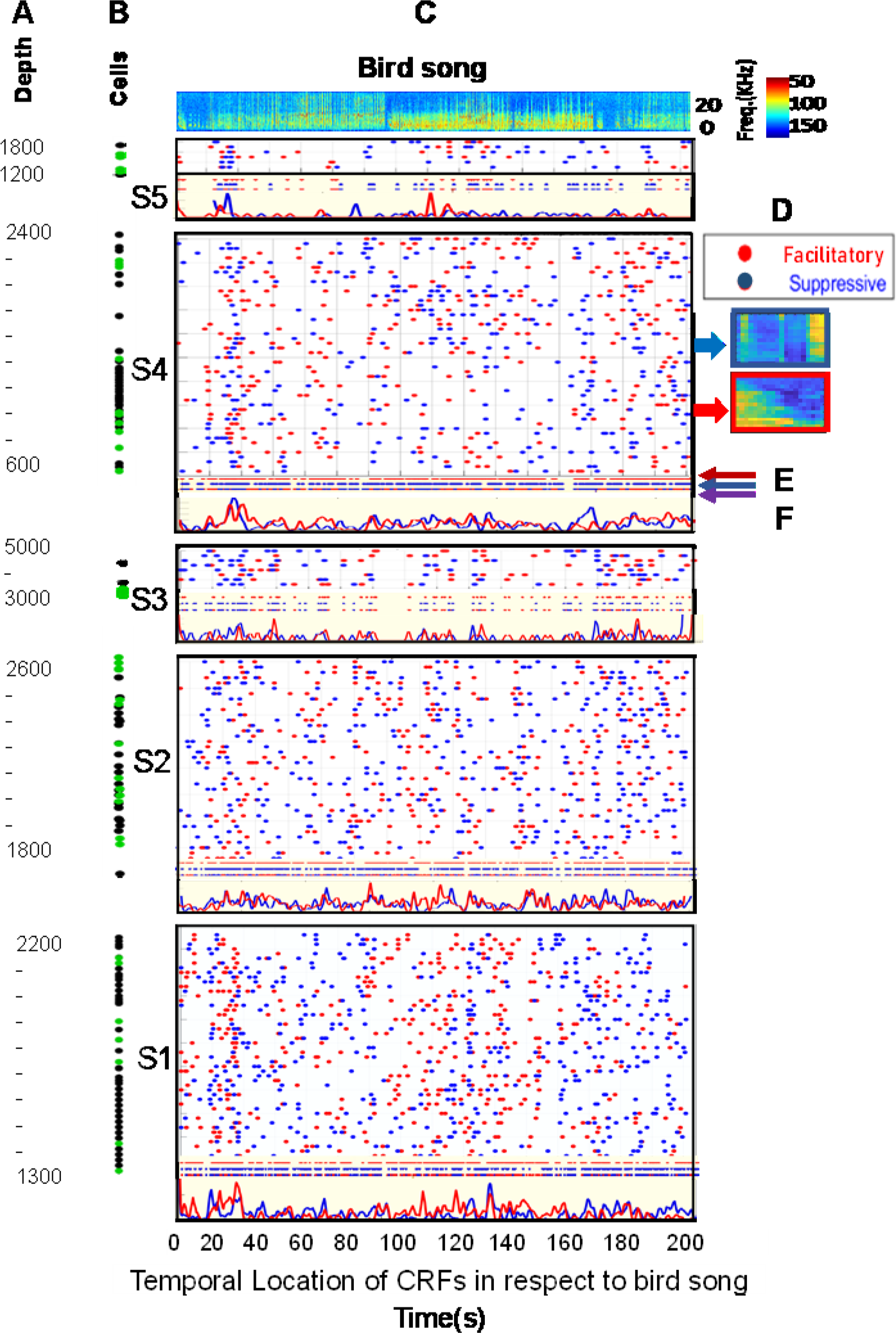
CRF Spatiotemporal activation maps. (A) Location of implanted electrodes in the caudomedial nidopallium for nine separate penetrations in five European starlings (S1-S5). (B) Spatial locations of the 154 single cells (green dots) and multi-unit clusters (black dots) used to create this map. The numbers of cells used from subjects S1-S5 were 50, 40, 8, 50, and 6, respectively. (C) Spectrogram of three concatenated bird songs. (D) Temporal location of CRFs with respect to bird song stimuli. There are 1540 facilitatory (red dots) and 1540 suppressive (blue dots) CRFs across the 154 cells in this map. Each dot corresponds to the maximum correlation (peak activation with r>60%) between each CRF and the song spectrograms. (E) Projections of all the facilitatory CRFs (red points), suppressive CRFs (blue line), and their joint (red and blue) CRFs onto the x-axis for every 20 bins. (F) Histogram of CRFs across cells for each subject (bin size=20). The red curves correspond to facilitatory CRFs, and the blue curves correspond to suppressive CRFs.

**Figure 4:**
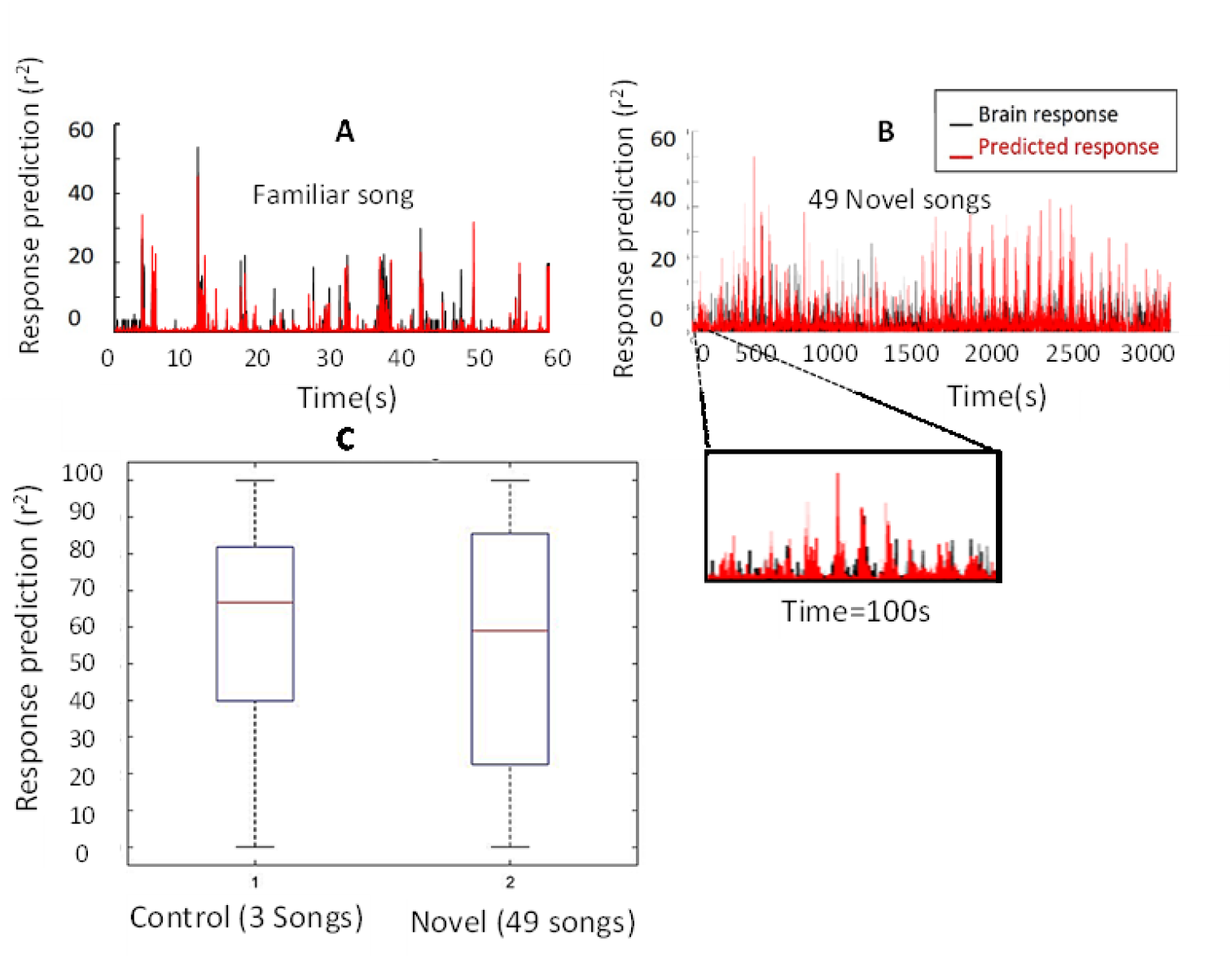
CRF response predictions. (A) Predicted response of a single neuron to a familiar song stimulus. Responses were predicted for five trials after training the model using the remaining 15 trials. The black curve shows the measured cellular response and the red curve shows the predicted response. (B) Predicted response of a single cell to 49 novel songs, where the model was trained on 48 songs and tested on the remaining song. The black curve shows the measured cellular responses to these 49 songs, and the red curve shows the predicted responses. All 49 songs are concatenated in this figure. The bottom panel shows the measured and predicted cellular responses for the first 100 s of songs. (C) Accuracy of the predicted response to three known songs (control test) compared with the accuracy of the predicted response to 49 novel songs. Boxplots show the results across ten different cells. The predicted response to the three familiar songs had a median of 65% accuracy, and the predicted response to 49 novel songs had a median of 60% accuracy, with a wider range than the control test.

We then mapped our population of neural cells, and their significant facilitatory and suppressive CRF features, to the location of implanted electrodes (i.e., a spatial map) and to the auditory stimuli time (i.e., a temporal map) (Figure 3). The spatial map was based on the location of each cell relative to an electrode channel. The location of each cell along the length of the electrode was calculated based on the reverse amplitude of the cell’s action potential (Rossant et al., 2016). On the other hand, temporal map for each cell and its CRFs, with respect to bird song stimuli, were characterized by correlating CRFs with the bird song spectrograms (Figure S2). Our spatiotemporal map of CRFs revealed several useful facts about the nature of CRFs. For example, the spatial map showed that this encoding mechanism for auditory stimuli is independent of neuron topography and location, whereas the temporal map suggested that neural processing of natural auditory stimuli follows a non-uniform distribution (Figures 5, 6).

**Figure 5:**
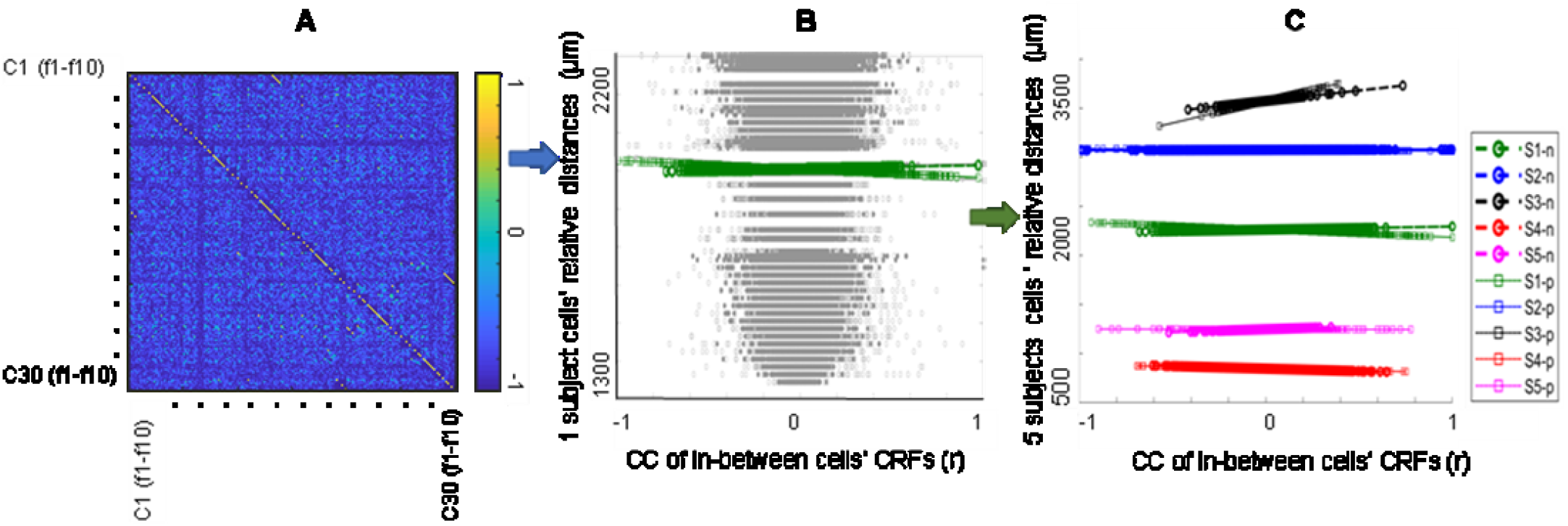
Spatial distribution of composite receptive fields (CRFs). (A) Example cross-correlation matrix for 10 facilitatory CRF features from 50 neurons in one subject (Subject S1). (B) Correlation coefficients (r) of CRFs between the 50 cells (excluding cells along the diagonal in panel [A]) versus the 50 cells relative distances for subject S1 for both facilitatory and suppressive responses in dark gray and light gray dots. The regression lines for facilitatory and suppressive CRFs are shown at the top using green circles and green squares, respectively. (C) Fitted regression lines for both facilitatory and suppressive CRFs from all five subjects. Each of the five subjects is identified with a unique color, and facilitatory (n) and suppressive (p) CRFs are distinguished using circled and squared lines, respectively.

**Figure 6:**
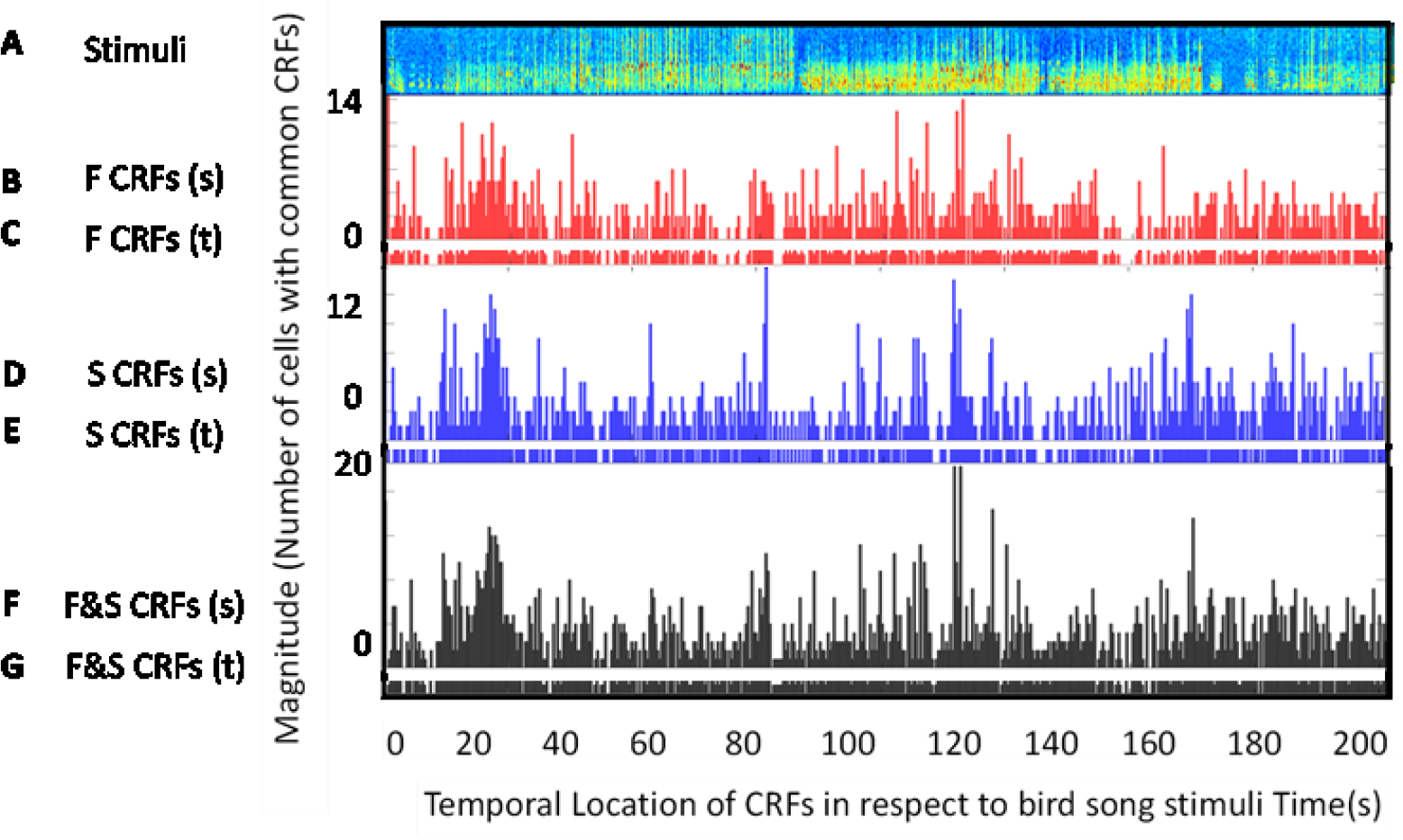
Temporal distributions of composite receptive field (CRF) features with respect to bird song stimuli. (A) Spectrogram of bird song stimuli. (B) Histogram of facilitatory CRFs for five subjects (S1 to S5) (C) The facilitatory histogram projected onto the x-axis. (D) Histogram of suppressive CRFs for five subjects. (E) The suppressive histogram projected onto the x-axis. (F) Sum of the facilitatory and suppressive CRF histograms. (G) The sum histogram projected onto the x-axis. The height of each histogram bar shows temporally co-active features that are common across sites.

One advantage of using CRFs generated from the MNE quadratic model versus receptive fields generated by linear models is that CRFs capture more variation in natural stimuli. As a result, their spectrograms are visually more similar to the spectrograms of bird songs (Figure 7B–F) (Ganji et al., 2019; Vahidi, 2019). Another advantage of CRFs over LRFs is that multiple CRFs can be extracted for each cell’s response, whereas LRFs are limited to one feature per cell.

**Figure 7:**
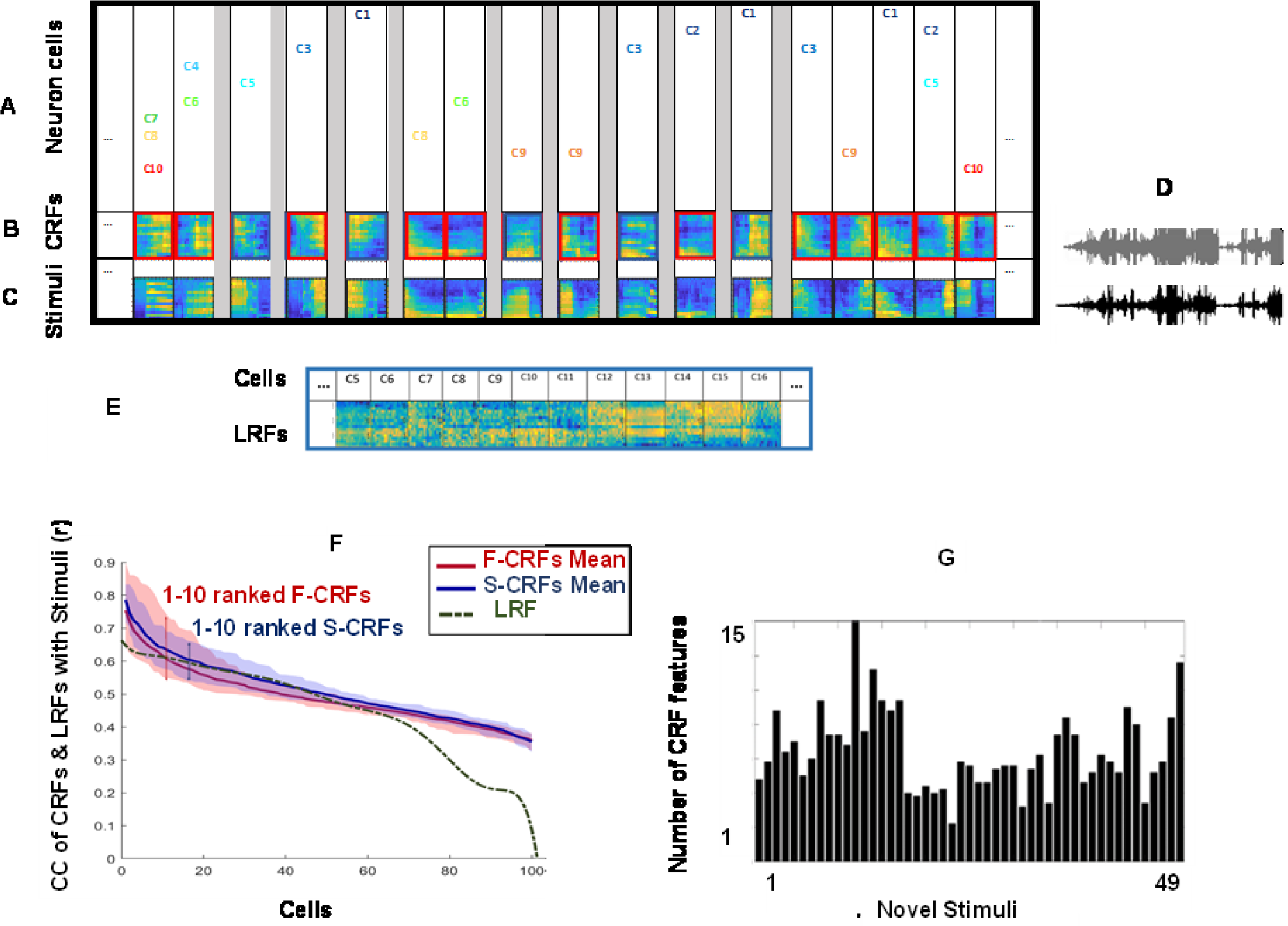
Stimuli encoding and rebuilding by CRFs. (A) A snap shot of connectivity map among populations of 10 cells. Some CRFs are shared among more cells (e.g. C7, C8, and C10 in second column) while some CRFs are shared among less cells (e.g. C5 in forth column). (B) Temporally organized CRF features. Facilitatory CRFs are shown by red frames and suppressive CRFs are shown by blue frames. (C) Stimuli rebuilding: Stimuli portions corresponding to each individual CRFs are found by the method from Figure S3 and stitched together. Gray spots will be further populated as the population of cells grow. (D) Bird song waveforms are extracted and reconstructed from both stitched CRFs (gray) and their corresponding stitched stimuli spectrograms (black). (E) Linear receptive fields (LRFs) of 12 cells extracted from the first order MNE model (h parameter). (F) Correlation coefficients between 100 cells facilitatory (red band) and supprasive CRFs (blue band) across CRFs with 1 to 10 ranks. Red and blue lines each indicates mean of 1000 CRFs (100 cells * 10 ranks). Cross-correlation coefficient (CC) of 100 LRFs (for 100 cells) with stimuli has shown in doted dark green curve. (G) Reconstructing of novel stimuli from pool of existing unique CRFs. Barographs display number of existing CRFs in each pieces of 49 equal length novel song (trained on 3 familiar songs and tested on each novel song at a time).

By taking advantage of these unique properties of CRFs, we have, for the first time, encoded and reconstructed both familiar and novel bird song stimuli from a pool of CRFs that we generated (Figure 7C, G). We additionally used our spatiotemporal CRF map to create a map of cell connectivity in response to stimuli by searching for common CRFs among different sub-groups of cells (Figure 7A). Our connectivity map highlights the fact that different sub-groups of neurons place varying emphasis on different parts of the stimuli (McGann, 2015). We further observed that the connectivity among different sub-groups of cells is altering by changes in the stimuli, indicating plasticity in the cellular response (Ho et al., 2011). Moreover, by projecting histogram bars from CRFs onto the temporal CRF map (x-axis), we discovered the percentage of a stimulus that was encoded by a certain number of cells, which allowed us to predict the number of cells and CRFs needed to encode an entire stimulus (Figure 8).

**Figure 8:**
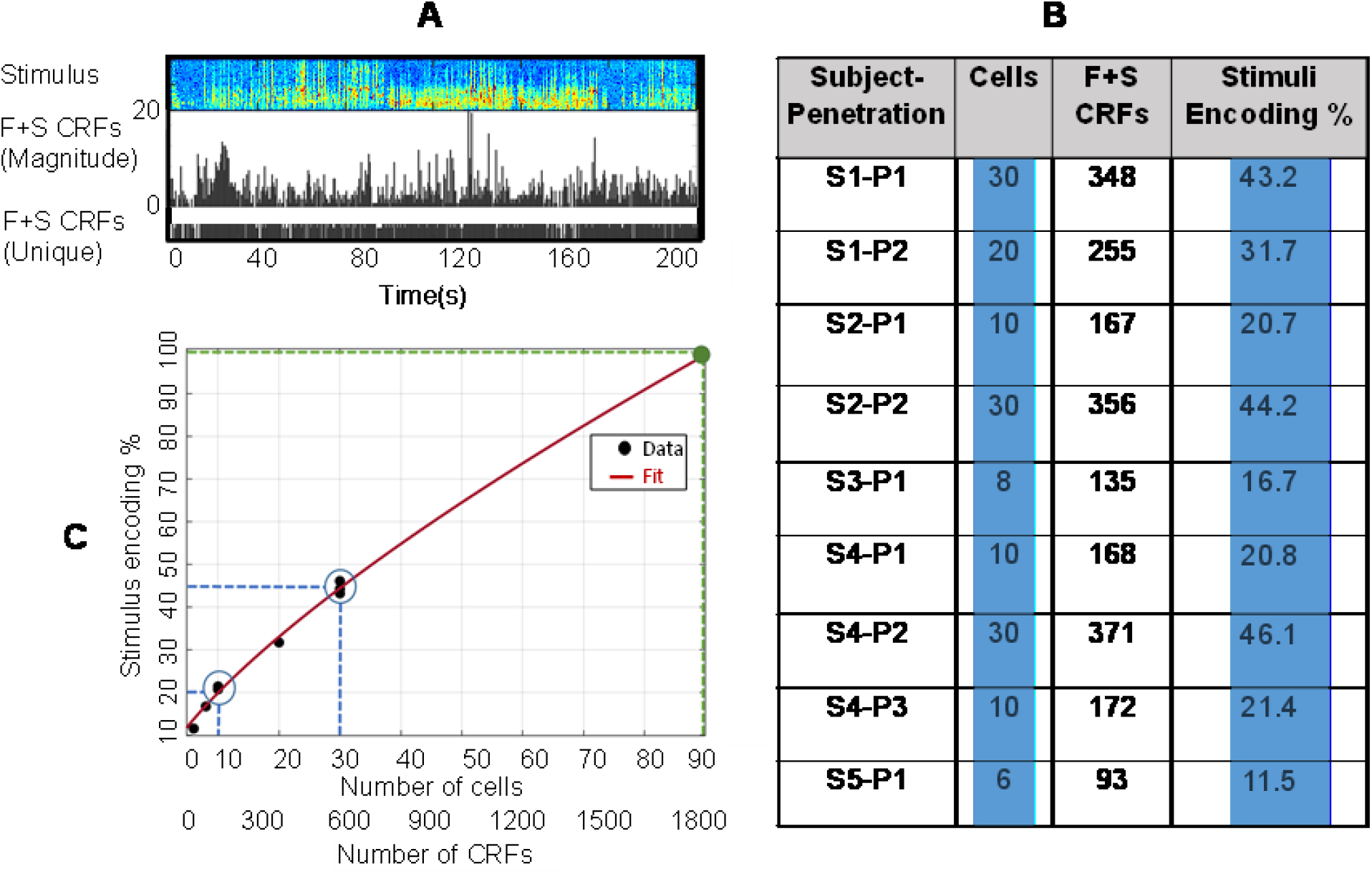
Prediction of number of cells and CRFs needed to encode entire stimuli. (A)First row: Spectrogram of stimuli. Second row: Sum of facilitatory and supprasive CRFs histograms and their flat temporal distribution projected onto x-axis in respect to 200s stimuli (See Figure 6). The bars flat projection onto x-axis has been shown underneath of the histogram in black dots. (B) First column: Nine recordings/penetrations across five subjects. Second column: Number of cells for each penetration. Third column: Sum of unique facilitatory and suppressive CRFs adopted from black dots on x-axis of figure A divided by (cells number*20 CRFs). Forth column (Stimuli encoding): Product of column three divided by column two. (C) Demonstration of relationship between cells numbers adopted from nine recordings (figure B second column) versus their stimuli encoding percentage (figure B fifth column). The red dotted line is a power fit and has been used to predict number of cells needed to encode entire stimuli (green dot). The predicated green dot here points to 90 cells as the required quantity of cells needed to encode this stimuli (contains three bird songs=207s). Out of the two blue circles marked here, one is pointing to three recordings with 10 cells and the other is pointing to three recordings with 30 cells that correspond to similar encoding percentages.

Overall, these outcomes demonstrate that CRFs can be used as a tool for mapping brain activity, encoding stimuli, and decoding neural activity in neuron populations. We can therefore use CRFs to improve our understanding of neural coding mechanisms in the brain.

## 2 Methods

All procedures were conducted in accordance with the guidelines of the Institutional Animal Care and Use Committee at the University of California, San Diego (S05383).

### 2.1 Animal subjects and surgical preparation

We recorded extracellular waveforms from the NCM of five lightly anesthetized (20% urethane, 7.0 ml/kg), adult male, wild-caught European starlings (mean weight: 90 g ± 0.9). Prior to the recording experiment, subjects were housed in large, mixed-sex outdoor aviaries with *ad libitum* access to food and water. On the day of the recording, subjects were anesthetized and placed in a stereotaxic holder. A fixation pin was glued to the top layer of the skull using dental acrylic. The top layer of the skull, dorsal to the electrode penetration site, was cleared away, and the subject was then transferred to the electrophysiological recording apparatus.

### 2.2 Electrophysiology

Each subject was placed on a foam bed in the recording apparatus, which was located inside a sound attenuation chamber (IAC Acoustics). The subject’s head was immobilized by the fixation pin. We first opened a small window in the lower skull (over the NCM); the window was only large enough to admit the recording electrode. We ruptured the exposed dura by making a small incision and then covered the dura with a silicone barrier. A subdural grounding wire was positioned several millimeters anterior to the electrode window. We lowered a 32-channel silicon electrode (NeuroNexus, A1×32-Edge-10mm-20-177) perpendicular to the exposed dorsal surface of the brain. For all recordings presented here, the distal tip of the recording electrode was positioned in a volume of tissue located 20-2500 μm anterior and 500-1700 μm lateral to the Y-sinus, at a depth of 1100-3200 μm. Table S1 indicates the exact position of the distal tip of the probe in each subject during the recordings. Continuous extracellular voltage waveforms from all electrode sites were passed through an analog headstage, bandpass filtered (300 Hz-10 kHz) and amplified (A-M systems model 3600), digitized with a 20 kHz sample rate (CED, power 1401, Spike2), and then stored for offline analysis. Subjects were euthanized at the end of the recording session, and brains were extracted for postmortem processing and to verify electrode placement within the NCM (Figure 1B).

### 2.3 Auditory stimulation

While the electrode was positioned at each recording site, we played three to five starling songs a minimum of 20 times each. Each song was roughly one minute long (66.6 s ± 2.93) and was recorded (44.1 kHz sample rate, 16 bit) from a single adult male that was unfamiliar to the subject. For some recording sites, we also presented 49 other starling songs that were recorded (44.1kHz, 16 bit) from both males and females. These songs were played once each. All songs were played from a small, full-range (20 Hz-20 kHz) speaker located inside the sound attenuation chamber. The mean sound pressure level was 60 dB (20 milipascals), measured at the approximate center of the animal’s head. Each auditory stimulus was digitized synchronously with the contemporaneous extracellular voltage waveforms (CED power 1401). Table S1 lists the stimuli presented at each recording location.

### 2.4 Composite receptive field estimation

We extracted clusters of similarly-shaped putative action potentials from the continuous extracellular voltage waveforms using the MountainSort (Chung et al., 2017) spike sorting package. We disregarded any spike clusters dominated by noise and attributed the action potentials in each remaining cluster to the response of either a putative single neuron or of multiple neurons, based on violations of a 2-ms refractory period (Dayan and Abbott, 2001). Table S1 indicates the numbers of single- and multi-unit clusters measured at each recording location.

We computed the CRF for each single- and multi-unit cluster using the MNE method (Fitzgerald et al., 2011a), which yields a minimally biased linear and nonlinear model for the stimulus-response functions of spiking neurons. The MNE method can identify multiple receptive-field features that are tied to the covariance structure of the stimulus, but unlike the STC method (de Ruyter van Steveninck et al., 1988), MNE works well with natural (i.e., non-Gaussian) stimuli. Specifically, the minimal model uses a logistic function to describe the probability of a spike given a stimulus *s*, as shown below:

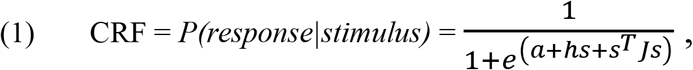

In this equation, the parameters *a*, *h*, and *J* represent the mean firing rate and the correlations with the first and second moments of the stimulus, respectively (Fitzgerald et al., 2011a). For our analyses, we down-sampled the stimuli to 24 kHz, converted them into log-scaled spectrograms (*nfft* = 128, Hanning window of length 128, and a 50% segment overlap), and removed the DC component following our previously published methods (Kozlov and Gentner, 2016). To reduce the dimensionality of the spectrographic stimulus representation, we averaged the power in pairs of adjacent frequency bands twice, which resulted in 16 frequency bands ranging from 750 Hz to 12 kHz. We then averaged adjacent time bins three times to obtain a final bin size of 21ms. To compute the CRF, we used 20 time-bins of the 16-channel spectrogram (320 dimensions total) as the stimulus, *s*, in Equation (1). We obtained similar CRFs when we used smaller time bins, a different number of 21ms time bins (10, 16, or 32), or 32 frequency bands for the stimuli dimensions (instead of 16) (see Kozlov and Gentner, 2016).

We determined CRF parameters by averaging the results from four separate estimates obtained using the responses in four disjoint subsamples. Each subsample comprised 25% of the total stimulus repetitions. For each of the four estimates, the data from the disjoint subsample was divided into training and test sets that comprised 75% and 25% of the data, respectively. We used early stopping for regularization to prevent overfitting. As in STC, diagonalizing the J-matrix yields quadratic features (eigenvectors) with the same time-by-frequency dimensions (20 × 16) as the original stimuli that evoked spiking. To avoid cases where a spurious response on one of the estimation runs could corrupt the average J-matrix, we clustered the t-SNE representation (van der Maaten and Hinton, 2008) of the J-matrix eigenvectors from all four estimation runs using the unsupervised HDBSCAN method (Ester et al., 1996). This method allowed us to objectively label and exclude any feature estimates that were extreme outliers (Figure S1B).

Overall, the negative eigenvectors of *J* correspond to facilitatory quadratic features of the CRF (i.e., those associated with relative increases in spike rates), and positive eigenvectors correspond to suppressive CRF features, associated with relative decreases in spike rates. The magnitude of the corresponding eigenvalues can be used to gauge the relative significance of each quadratic feature of the CRF (Figure 2B). To set a criterion value, we computed the 95 percent confidence interval around the distribution of eigenvalues from 500 symmetrical random Gaussian matrices with size and moments equal to J. Empirical eigenvalues outside this interval are taken to be significant. Eigenvalues closer to zero correspond to noise or to features whose contribution to the response is beyond the sensitivity of our MNE method. To minimize noise, and ease comparisons across units, we selected the ten most significant positive and negative quadratic features (20 total) from each CRF. This accords roughly with the number of significant features relative to the 95% CI, though likely includes some noisy features in some CRFs. Figure 2C shows an example of CRFs extracted from a single cell with the ten most significant negative (facilitatory) and ten most significant positive (suppressive) features.

### 2.5 CRF response predictions

To test how well the CRF models predict responses to a given stimulus, we computed the CRF parameters a, h, and J for each neuron using n-1 of the n stimuli presented at a given site (as described above), with the target (to-be-predicted) stimulus and response held out (Figure 2A and Figure S1A). Substituting the estimated parameters and the target stimulus into equation (1), we obtained a predicted response to the target stimulus. In practice, low-rank reconstructions of the J matrix (rank=20) yield better SNR for response predictions than those obtained from the full rank matrix (rank=320) (Kaardal et al., 2017). For the predictions reported here, we reconstruct J from only the first 10 facilitatory and first 10 suppressive features (rank = 20) using singular value decomposition (SVD). We compare the predicted response to the empirical response for each stimulus using the Matlab corrcoef function. For a subset of units, we predicted the neural response of unique repetitions of the same stimuli used to estimate the CRF, and for other units on a large set (n=49) novel song stimuli ranging in length from 30 to 90 sec (Figure 4 A, B).

### 2.6 Relationship between CRFs and stimuli (Spatial-Temporal map of CRF)

To investigate this relationship, we have generated a spatial and temporal map of cells’ CRF responses to their stimuli. To compute the cells’ CRFs temporal map, a normalized cross-correlation method is used. This model cross-correlates the power spectrogram of each CRFs with the power spectrogram of stimuli through a sliding window to locate the maximum cross-correlation of each CRFs with the stimuli (Figure 3). Ultimately, these maximum correlations (peak activations) are tested to assure more than a 60% correlation between CRFs and Stimuli.

Figure S2C displays how to find the temporal location of 20 facilitatory and suppressive CRFs of one cell corresponding to specific portions of vocal elements of stimuli. Figure S2A demonstrates the time bins that each CRFs corresponds to its stimuli portion.

To build the spatial map, cells are organized based on their locations along the dorsal-ventral plane (z-axis) in the NCM auditory area. The spatial (depth location) information of cells is calculated via the Klustakwik program (Rossant et al., 2016). The cell locations are calculated based on their inverse action potential amplitude of each cell in reference to the z-axis (depth) of the electrode shank.

Finally, the above two methods are combined to investigate the spatial-temporal map of CRFs of the population of cells with respect to stimuli. To generate this map, we have extracted CRFs from 154 cells. These cells are chosen from nine recording sessions across five subjects (Table S1). Then ten most negative (facilitatory) and ten most positive (suppressive) CRF features of each of these cells are extracted to create a pool of 3080 CRFS ((10f +10s)*154 cells). Figure 3 demonstrates a spatial-temporal map of the pool of the 154 cells and their 3080 CRFs. Figure 3A displays the locations of neural cell recordings along the dorsoventral plane in the NCM for nine penetrations across five subjects. Next to each penetration, the recorded cells are shown which are organized spatially by their depth locations. The cells are either single cells in green or multi-units in black (Figure 3B). Figure 3C displays a spectrogram of three concatenated bird songs. Underneath, Figure 3D displays temporal maps of 3080 CRFs extracted from 154 cells with respect to the stimuli across five subjects. The facilitatory CRFs are shown in red dots (3080/2=1540) and the 1540 suppressive CRFs are shown in blue dots. Each dot corresponds to the maximum correlation (peak activation) of each CRF with the spectrogram of songs. The temporal mapping can be utilized as a method to encode stimuli by CRFs. Figure 3E demonstrates projections of facilitatory (in red dots), suppressive (in blue dots), and their joint (red and blue dots) on temporal x-axis. This demonstrates that even a small sample, e.g. 50 cells (from subject 1), can provide a temporally dense representation of the natural song. To investigate the temporal distribution of CRFs with more details, we have created histograms of the temporal distribution of CRFs of cells for all subjects (e.g. Figure 3F for subject 2). Red curves correspond to facilitatory CRFs and blue curves correspond to suppressive CRFs with bin sizes of 20.

Overall, this map not only can be used to demonstrate the spatial and temporal relationship between the population of cells’ CRFs and stimuli but also, we will show in the following sections, this map can aid us to investigate questions such as how CRFs are distributed across time and space, can stimuli be encoded and reconstructed by CRFs, and if the answer is affirmative, then how many cells and CRFs are needed to encode entire stimuli?

## 3 Results

We recorded the song-evoked extracellular spiking responses of 40 single-unit and 115 multi-unit neurons obtained from nine separate penetrations into the NCM in five adult European starlings (Table S1). Responses were recorded simultaneously in neural populations that ranged in size from 6 to 30 units (Figure 1) and were detected using 32-channel linear arrays that were oriented along the dorsal-ventral axis of the NCM. For each isolated single unit and multi-unit cluster, we computed the CRF (Figure 2), which provides an estimate of the probability of a response given a stimulus. Each individual CRF is an estimate of the linear and nonlinear components that comprise that unit’s receptive field. These linear and non-linear components can be abstractly considered to be the classic STA and STC, respectively, or can be viewed more concretely as acoustic “features” of the training stimuli that evoke or suppress spiking responses. These acoustic features may reflect simple, general properties of the stimulus, such as the power of the stimulus in a specific frequency range, or they may reflect spectro-temporal events that are more complex (and rare).

The observation of a CRF implies that the magnitude of the response in a given neural unit cannot be interpreted as a stationary scalar estimate of the likelihood of a single stimulus event or feature. Instead, changes in a unit’s response at different points in time may correspond to the presence (or absence) of different stimulus features. This interpretation agrees with our qualitative observations of neural responses in the NCM (Figure 1C, D), where single-unit responses were tightly time-locked to multiple seemingly distinct features that were distributed throughout the duration of conspecific songs. Our overall goal was to understand how the features of CRFs for individual units are spatially organized within the NCM and how these features correspond to changes in activity over the duration of natural conspecific communication signals (songs).

### 3.1 Predictive quality of the CRFs

Before examining the organization of CRF features in detail, we first confirmed that the CRFs provided a reasonable explanatory model for the response characteristics of a given unit. In general, the MNE method that we used to compute CRFs produced significantly better predictions of neural responses to natural stimuli than comparable linear techniques (Kozlov and Gentner, 2016). We used Eq. (2) to predict the neural response to stimuli:

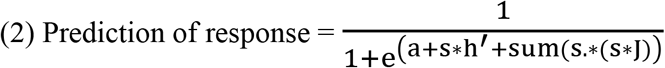

In this equation, *s* is the stimulus and *a, h*, and *J* represent the constant, linear, and quadratic parameters, respectively, of the receptive fields. To evaluate this equation, we first ran a control test using a cell for which we recorded the response to three familiar bird songs across 20 trials. Fifteen of these trials were used to train the equation and estimate the parameters *a*, *h*, and *J*. The estimated parameters were then used to predict the responses for the five remaining trials. The black curve in Figure 4A shows the average measured brain response to one 60-second bird song and the red curve shows the predicted response estimated using Eq. 2.

We repeated this test of Eq. 2 using ten additional cells. The results for these additional cells are shown in Figure 4C. The box plot on the left side of Figure 4C shows that the predicted response had a 40-80% correlation with the measured response (median 65%, red bar), indicating that our proposed prediction model (Eq. 2) was reliable.

After confirming the reliability of our prediction equation, we predicted brain responses to 49 novel stimuli using a single cell for which we measured responses to 49 novel songs over 49 trials. We first trained the equation and estimated the parameters *a, h*, and *J* using data from 48 of these trials. The estimated parameters were then used to predict the response for the remaining trial (Eq. 3).

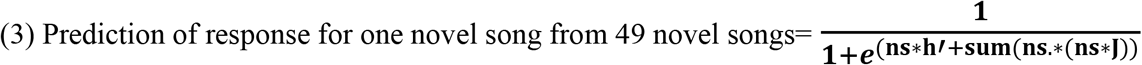

In Equation 3, *ns* represents a novel song

All 49 responses measured for the cell were estimated using this same procedure. Figure 4B shows the 49 predicted responses for the cell (red curve) relative to the 49 observed responses for the cell (black curve). We repeated this test using data for ten additional cells. The box plot on the right of Figure 4C shows the accuracy of predicted responses to 49 novel songs across these ten cells. Although the prediction range varied from 25% to 85%, the prediction median was about 60%. Based on the results of our model for both three familiar songs (control) and 49 novel songs, the MNE model is a good candidate for generating receptive fields using data obtained from neural cells. These results also indicate that we can accurately predict brain responses using CRFs.

### 3.2 Spatial and temporal distribution of CRFs

To evaluate the spatial distribution of CRFs, we tested for similarity in the CRFs of cells taken from each subject using the autocorrelation method. Figure 5A provides an example of an autocorrelation matrix for 50 cells from Subject 1. In this example, each cell contains 10 facilitatory CRFs. Correlation coefficients (r) were extracted from the autocorrelation matrix (except for the cells along the diagonal) to evaluate the similarity of CRFs between cells. Figure 5B provides an example of between-cell CRF similarity of 50 cells for Subject 1versus these 50 cells relative physical distances for both facilitatory and suppressive responses in dark gray and light gray dots. The regression lines for facilitatory and suppressive CRFs are shown at the top using green circles and green squares, respectively.

We repeated this process to extract between-cell CRF similarity information for both facilitatory and suppressive responses from each of the five subjects. We then calculated relative distances between cells for each subject. The fitted regression lines for both facilitatory and suppressive responses of all subjects are shown in Figure 5C. Subjects 1 to 5 contained 50, 40, 6, 50, and 8 cells, respectively. Each of the five subjects shown in Figure 5C is identified with a unique color, and facilitatory (n) and suppressive (p) CRFs are distinguished using circled and squared lines, respectively. The distance between circles in the facilitatory regression lines and between squares in suppressive regression lines indicates the CRF similarity density, which is high around r=0 and decreases toward r=1 or −1. Most of the regression line slopes for both facilitatory and suppressive CRFs are constant around different y-intercepts (distances). There is a slight slope for Subject 3 (black regression lines) which could be because cells from Subject 3 contained more noise than cells from other subjects. Overall, the low slopes of these regression lines indicate that between-cell CRF similarity for both facilitatory and suppressive responses across all subjects is independent of the relative distances between cells along the dorsoventral plane. In other words, stimuli encoding by neuron populations is independent of the locations and topology of the neurons.

To assess the temporal distribution of CRFs, we used the histograms of 3,080 CRFs for all five subjects shown in our spatiotemporal map (Figure 3F). Figure 6 displays these histograms and their projects on x-axis with respect to the auditory stimuli. The height of each bar in the histogram shows temporally co-active features that are common across cells. The histogram with red bars represents facilitatory CRFs (Figure 6B), and the histogram with blue bars represents suppressive CRFs (Figure 6D). Figure 6F shows the sum of facilitatory and suppressive CRFs. Underneath each histogram (Figure 6C, E, and G) projection of the histogram bars onto the x-axis are shown as points.

The overall temporal distribution of CRFs was obtained by taking one hundred samples one hundred times from these points. Samples were tested using a one-sample Kolmogorov-Smirnov test with a significance criterion of p < 0.05. Our results reject the hypothesis that the temporal distribution of CRFs follows a standard normal distribution.

In conclusion, the spatial map showed the CRFs of neurons encode auditory stimuli independent of neurons location and topology while the temporal map suggested that neural processing of auditory stimuli follows a non-uniform distribution.

### 3.3 Stimuli encoding and reconstructing by CRFs

Using our spatiotemporal map of CRFs, we were able to encode the various stimuli by connecting peak activation of receptive fields to corresponding portions of the stimuli. In this section, we demonstrate this encoding mechanism. Figure S2 provides an example of the method we used to find portions of a 200-second stimulus that were associated with 20 CRFs. Figure S2B shows the 20 CRF features, 10 facilitatory (red frames) and 10 suppressive (blue frames), that were adopted from one cell. The temporal locations of these CRFs, which were determined based on their peak activation relative to the stimuli power spectrogram, are shown in Figures S2C and S2A (r>60%). We found the portions of the stimuli that correspond to the CRFs by referring to the temporal locations within the stimuli (Figure S2D).

When we concatenated these CRFs based on their temporal organization, we produced Figure 7B. By identifying the peak activation of these CRFs in relation to the stimuli spectrograms, as shown in Figure S2, we can find the time window of the stimulus that corresponds to CRF activation. Figure 7C displays these corresponding sections of the stimuli arranged in temporal order. Figure 7D demonstrates the powers extracted from stitched CRFs as well as the stimuli spectrograms of Figures 7B and C, which were then reconstructed back to audio waveforms.

Based on the results shown in Figure 7, we concluded that the concatenated CRFs contain a reasonable amount of information about the corresponding portions of the stimuli. Such a large amount of information is obtainable because we captured the receptive field responses using the quadratic MNE model rather than LRFs. Figure 7E displays the LRFs of 12 cells from the second penetration in Subject 2. These LRFs are extracted from the linear parameter of the MNE model (*h* in Eq. 1).

To compare how similar CRFs and LRFs were to the stimuli spectrograms, we ran cross-correlations between 200-second stimuli and 100 cells that each contained ten rank-ordered facilitatory CRFs (i.e., the first ten CRFs out of 320), ten rank-ordered suppressive CRFs, and one LRF. The results are shown in Figure 7F. The top 100 facilitatory and the top 100 suppressive CRFs across all ten ranks were more highly correlated with the stimuli than the top 100 LRFs. Quadratic CRFs therefore appear to contain more information about the stimuli than LRFs and could potentially be used to rebuild and reconstruct stimuli. CRFs can also be extracted in large numbers from each cell, whereas LRFs are limited to one feature per cell.

The fact that we were able to successfully reconstruct parts of the 200 s stimulus (which contained three concatenated stimuli) using a small group of CRFs encouraged us to test the hypothesis that the total pool of 3,080 CRFs could be used to rebuild portions of novel stimuli. For this test, we acquired the 49 novel songs that were played for the subjects during the brain response recordings. These songs were considered “novel” because they were only played for the subjects once and they were not tested in any of our previous analyses. The novel songs varied in length from 30 s to 90 s, so we concatenated all the songs and then divided them into 49 equal-length stimuli. We then extracted unique CRFs from the pool of existing CRFs. Unique CRFs were the points generated by the projection of histogram bars onto the x-axis and therefore do not encompass recurrent CRFs (Figure 6C, E, G). Of the 3,080 CRFs that we pooled from 154 cells, 646 were unique and 2,434 were recurrent. The spectrogram of each piece of the 49 songs was then cross-correlated with the 646 unique CRFs and then the unique CRFs with correlation coefficients larger than 60% with novel songs were saved. Each bar in Figure 7G displays the number of these unique CRF features that existed in all 49 novel songs. Ultimately CRFs and their corresponding stimuli portions can be extracted from each bar and reconstructed in the form of bird song waves (similar to Figure 7Din conclusion, we showed that not only can parts of bird song be rebuilt using the pool of CRFs, but novel portions of bird song can also be reconstructed using the existing CRFs. To improve the reconstruction accuracy, CRFs should be extracted from a larger training set (i.e., with longer birdsongs) and then tested on each novel song one at a time.

### 3.4 Map of cell connectivity using CRFs

We additionally used our spatiotemporal CRF map to evaluate connectivity among cells. Figure 7A provides an example subset of the “connectivity map” that were generated among populations of 10 cells. We observed that some CRFs are shared across more cells (e.g., C7, C8, and C10 in the second column) while other CRFs are shared across fewer cells (e.g., C5 in the fourth column). Gray spots will be further populated as the population of cells grows. These observations emphasize the fact that sub-populations of cells encode stimuli intensity by placing different weights on different parts of the stimuli. This means the connectivity map among cells changes with changes in the stimuli, which can be indications of cell response flexibility (Hubel and Wiesel, 1959). This fact might save energy and time for the brain during stimulus encoding (Rehn and Sommer, 2007).

### 3.5 Prediction of number of cells and CRFs needed to encode entire stimuli

In Section 3.3, we showed that stimuli can be encoded and reconstructed using CRFs. The logical next question is how many CRFs and cells are needed to encode an entire stimulus. To answer this question, first we calculated the number of CRFs that encoded stimuli for each recording session. Because each CRF image has 20 bins, multiplying the number of CRFs by 20 and then dividing the product by the length of the stimuli gives the percentage of the stimulus that was encoded by the CRFs, which can then be used to calculate the number of cells needed to encode the entire stimulus. This process is explained in Figure 8. Figure 8A is adapted from the fourth row of Figure 6. The histogram bars represents the cumulative number of facilitatory and suppressive CRFs. The black points which are the result of projection of the histogram bars onto the x-axis is shown underneath the histogram.

Figure 8B shows how we calculated the percentage of the stimulus that was encoded in each of our nine recordings. The first column shows the identifiers of the nine recordings across five subjects, and the second column shows the number of cells per recording. In the third column, the sum of unique facilitatory and suppressive CRFs (determined from the black points in Figure 8A) divided by a default number of facilitatory and suppressive CRFs. This default number was determined by multiplying the number of cells by 20 CRFs (10 facilitatory and 10 suppressive) per cell. The last column is the value in the third column divided by the value in the second column, such that the value in the last column indicates the percentage of the stimulus that was encoded by the number of cells listed in the second column.

As an example, consider the first penetration in Subject 1 (S1-P1), which sampled 30 cells (first and second columns of Figure 8B). From the default pool of 600 CRFs for this recording (i.e., 30 cells * 20 CRFs = 600), there were 348 unique facilitatory and suppressive CRFs that temporally encoded the stimuli (see Figure 8A). Because each CRF has 20 time bins, all 348 CRFs together cover 6,960 time bins (i.e., 348 CRFs * 20 bins = 6,960 bins). When we divide this number by the length of the stimulus in this study (207 seconds, or 16,100 time bins), we find that 348 CRFs extracted from 30 cells encode 43.2% of the stimulus (Figure 8B, fourth column).

Figure 8C demonstrates the relationship between the number of cells measured in each penetration with the percentage of the stimulus that they encoded (adapted from the second and fourth columns, respectively, of Figure 8B). This figure shows that the number of cells is approximately linearly related to the percentage of the stimulus that they encode (red line).

The green dot in Figure 8C, located at the intersection of the dotted green lines from the x- and y-axes, represents the predicted number of cells needed for the CRFs to encode the entire stimulus. In this case, we predicted that 100% of the 207-second stimulus would be encoded by 90 cells and 1,800 CRFs (i.e., 90 cells * 20 CRFs per cell). It is worth noting that this prediction paradigm is only possible because our nine penetrations sampled cell populations of varying sizes. We additionally noted that, when using this calculation paradigm, choosing similarly-sized cell populations from different recordings resulted in similar stimuli encoding percentages; for example, the population of 10 cells in recordings S2-P1, S4-P1, and S4-P3 all led to a similar average encoding of approximately 20%.

The population of 30 cells in recordings S1-P1, S2-P1, and S4-P2 likewise corresponded to an average encoding of 45%. The high degree of agreement between the results of different penetrations emphasizes the accuracy of the CRF stimulus encoding mechanism. Our method could also be used to investigate the stimulus encoding percentage separately for facilitatory and suppressive CRFs.

## 4 Discussion

We previously showed that neurons in the secondary auditory cortical regions of European starling brains have CRFs (Kozlov and Gentner, 2016). In this study, we used CRFs from a population of neurons in the auditory forebrain of starlings to, for the first time, investigate a spatiotemporal map of CRFs and the associated properties of CRFs along the dorsoventral brain axis. To generate CRFs from neurons, we used the MNE quadratic model (Fitzgerald et al., 2011a). This model has been proven to be more effective than other common approaches such as STA, STC, and MID (de Ruyter van Steveninck et al., 1988; Sharpee et al., 2004; Bialek and de Ruyter van Steveninck, 2005). For example, the MNE model does not require the user to work with natural stimuli, and it can extract many CRF features from cellular responses (Sharpee et al., 2004; Kell et al., 2018). We further examined the validity of the MNE model and its neural response decoding mechanism by utilizing CRF features to predict neural responses to novel stimuli. Our results showed that neural responses can be accurately predicted and reconstructed using CRFs.

Neural response prediction has extensive applications for brain-computer interface research. For example, it can assist patients who have speech impairments caused by neurological disorders (Anumanchipalli et al., 2019; Pandarinath and Ali, 2019). By utilizing the MNE model, we created a large pool of facilitatory and suppressive CRFs from 154 auditory NCM cells sampled during nine recording sessions across five European starlings. Since high-quality CRFs generated from stimuli-locked cells with low noise are preferred, a combination of techniques is suggested to remove noisy trials from cells if required ((Ester et al., 1996; van der Maaten and Hinton, 2008).

To investigate the mechanisms by which CRFs encode stimuli, we used our spatiotemporal map of CRFs. The map was created from the assembled pool of CRFs. The spatial component of the map reflects the dorsal-ventral organization of our population of 154 cells, and the temporal component reflects the location of peak activation of each CRF with respect to a specific portion of the stimuli. As a result, the map shows the spatial and temporal distribution of CRFs as well as the stimuli encoding process. By investigating the spatial distribution of CRFs, we showed that the relative locations of cells along the dorsoventral brain axis are independent of any similarities in their responses, which suggests that stimuli encoding by a population of neurons is independent of neuron location or topology. Furthermore, temporally, CRFs appeared to follow a non-uniform distribution.

Another property of CRFs that we discovered using our spatiotemporal CRF map is their ability to practically encode and construct stimuli. We showed that CRFs carry enough information about their corresponding stimuli portions to reconstruct the stimuli. This is because we captured CRFs using the quadratic MNE model instead of a linear model. By monitoring peak activation of CRFs in relation to various stimuli, we were able to locate the corresponding spectrograms of the stimuli. For example, by concatenating the stimuli spectrograms and extracting their powers, we reconstructed parts of familiar bird songs and of various novel songs. To improve the stimuli reconstruction accuracy, CRFs should be extracted from a larger training set and be tested on short, novel songs.

In addition to revealing this novel encoding mechanism, our spatiotemporal CRF map revealed other remarkable properties of CRFs. For instance, by concentrating the projections of CRFs on histograms on the x-axis, we not only found the percentage of stimuli encoding for different cell populations but were also able to predict the number of cells and CRFs needed to encode entire stimuli.

We were further able to monitor cell connectivity and organization using our spatiotemporal CRF map. We observed that some CRFs are shared among more cells while others are shared across fewer cells. This observation highlights the fact that sub-populations of cells place different weights on different parts of stimuli, as evidenced by their responses. This process saves time and energy among a group of cells while still ensuring the input stimuli is encoded (Rehn and Sommer, 2007). In addition, our connectivity map shows that sub-populations of cells connect and rewire simultaneously with changes in the pattern and intensity of the stimuli, indicating plasticity in the cellular response (Ho et al., 2011).

It is worth noting that all the characteristics of CRFs described above were tested on both facilitatory and suppressive CRF responses, and both responses appear to have similar neural coding mechanisms.

Overall, our results demonstrate that CRFs can be used as tools for mapping brain activity, encoding stimuli, and decoding neural activity in neuron populations, which can improve our understanding of neural coding mechanisms in the brain. CRFs generated from neural responses to auditory stimuli also have practical applications; for example, they can be used to reconstruct human speech to design better hearing aids (Schäfer et al., 2018) or to create speech decoders to help speechless patients communicate (Rieke et al., 1995; Stanley et al., 1999; Zion Golumbic et al., 2013).

## 5 Conflicts of Interest

The author declare that the research was conducted in the absence of any commercial or financial relationships that could be construed as a potential conflict of interest.

## 6 Author Contributions

N.W.V. was involved in the implementation, programming, calculations, and manuscript writing.

## 7 Acknowledgments

I thank Dr. Timothy Gentner for suggesting study hypothesis and manuscript editing.

## 8 Funding

This work was supported by the Dynamics of Multifunction Brain Network (ONR) and the Rita L. Atkinson fellowship.

## Notes

### Competing Interest Statement

The authors have declared no competing interest.

## References

Anumanchipalli, G. K., Chartier, J., and Chang, E. F. (2019). Speech synthesis from neural decoding of spoken sentences. Nature 568, 493–498. doi:10.1038/s41586-019-1119-1.

Bialek, W., and de Ruyter van Steveninck, R. R. (2005). Features and dimensions: motion estimation in fly vision. arXiv. Available at: https://arxiv.org/abs/q-bio/0505003.

Chung, J. E., Magland, J. F., Barnett, A. H., Tolosa, V. M., Tooker, A. C., Lee, K. Y., et al. (2017). A fully automated approach to spike sorting. Neuron 95, 1381–1394.e6. doi:https://doi.org/10.1016/j.neuron.2017.08.030.

Dayan, P., and Abbott, L. E. (2001). Theoretical Neuroscience: Computational and Mathematical Modeling of Neural Systems. Cambridge, MA: Massachusetts Institute of Technology Press.

de Boer, R., and Kuyper, P. (1968). Triggered correlation. IEEE Trans. Biomed. Eng. 15, 169–179. doi:10.1109/tbme.1968.4502561.

de Ruyter van Steveninck, R., Bialek, W., and Barlow, H. B. (1988). Real-time performance of a movement-sensitive neuron in the blowfly visual system: coding and information transfer in short spike sequences. Proc. R. Soc. London. Ser. B. Biol. Sci. 234, 379–414. doi:10.1098/rspb.1988.0055.

Eggermont, J. J., Epping, W. J., and Aertsen, A. M. (1983). Stimulus dependent neural correlations in the auditory midbrain of the grassfrog (*Rana temporaria L.*). Biol. Cybern. 47, 103–117. doi:10.1007/BF00337084.

Ester, M., Kriegel, H.-P., Sander, J., and Xu, X. (1996). A density-based algorithm for discovering clusters in large spatial databases with noise. in Proceedings of the Second International Conference on Knowledge Discovery and Data Mining KDD’96. (AAAI Press), 226–231.

Fitzgerald, J. D., Rowekamp, R. J., Sincich, L. C., and Sharpee, T. O. (2011a). Second order dimensionality reduction using minimum and maximum mutual information models. PLoS Comput. Biol. 7, e1002249. Available at: https://doi.org/10.1371/journal.pcbi.1002249.

Fitzgerald, J. D., Sincich, L. C., and Sharpee, T. O. (2011b). Minimal models of multidimensional computations. PLoS Comput. Biol. 7, e1001111. Available at: https://doi.org/10.1371/journal.pcbi.1001111.

Ganji, M., Paulk, A. C., Yang, J. C., Vahidi, N. W., Lee, S. H., Liu, R., et al. (2019). Selective formation of porous Pt nanorods for highly electrochemically efficient neural electrode interfaces. Nano Lett. 19, 6244–6254. doi:10.1021/acs.nanolett.9b02296.

Harris, K. D. (2015). Cortical computation in mammals and birds. Proc. Natl. Acad. Sci. 112, 3184 LP – 3185. doi:10.1073/pnas.1502209112.

Hartline, H. K. (1938). The response of single optic nerve fibers of the vertebrate eye to illumination of the retina. Am. J. Physiol. 121, 400–415. doi:10.1152/ajplegacy.1938.121.2.400.

Ho, V. M., Lee, J.-A., and Martin, K. C. (2011). The cell biology of synaptic plasticity. Science 334, 623–628. doi:10.1126/science.1209236.

Hubel, D. H., and Wiesel, T. N. (1959). Receptive fields of single neurones in the cat’s striate cortex. J. Physiol. 148, 574–591. doi:10.1113/jphysiol.1959.sp006308.

Kaardal, J. T., Theunissen, F. E., and Sharpee, T. O. (2017). A low-rank method for characterizing high-level neural computations. Front. Comput. Neurosci. 11, 68. doi:10.3389/fncom.2017.00068.

Karten, H. J. (2013). Neocortical evolution: neuronal circuits arise independently of lamination. Curr. Biol. 23, R12–R15. doi:https://doi.org/10.1016/j.cub.2012.11.013.

Kell, A. J. E., Yamins, D. L. K., Shook, E. N., Norman-Haignere, S. V, and McDermott, J. H. (2018). A task-optimized neural network replicates human auditory behavior, predicts brain responses, and reveals a cortical processing hierarchy. Neuron 98, 630–644.e16. doi:10.1016/j.neuron.2018.03.044.

Koyama, S. (2012). On the relation between encoding and decoding of neuronal spikes. Neural Comput. 24, 1408–1425. doi:10.1162/NECO_a_00279.

Kozlov, A. S., and Gentner, T. Q. (2016). Central auditory neurons have composite receptive fields. Proc. Natl. Acad. Sci. 113, 1441 LP – 1446. doi:10.1073/pnas.1506903113.

McGann, J.P., (2015). Associative learning and sensory neuroplasticity: how does it happen and what is it good for? Learn Mem. 22(11):567–76. doi: 10.1101/lm.039636.115.

Mesgarani, N., David, S. V, Fritz, J. B., and Shamma, S. A. (2009). Influence of context and behavior on stimulus reconstruction from neural activity in primary auditory cortex. J. Neurophysiol. 102, 3329–3339. doi:10.1152/jn.91128.2008.

Pandarinath, C., and Ali, Y. H. (2019). Brain implants that let you speak your mind. Nature 568, 466–467. doi:10.1038/d41586-019-01181-y.

Pasley, B. N., David, S. V, Mesgarani, N., Flinker, A., Shamma, S. A., Crone, N. E., et al. (2012). Reconstructing speech from human auditory cortex. PLOS Biol. 10, e1001251. Available at: https://doi.org/10.1371/journal.pbio.1001251.

Quiroga, R. Q., Reddy, L., Kreiman, G., Koch, C., and Fried, I. (2005). Invariant visual representation by single neurons in the human brain. Nature 435, 1102–1107. doi:10.1038/nature03687.

Rehn, M., and Sommer, F. T. (2007). A network that uses few active neurones to code visual input predicts the diverse shapes of cortical receptive fields. J. Comput. Neurosci. 22, 135–146. doi:10.1007/s10827-006-0003-9.

Rieke, F., Bodnar, D. A., and Bialek, W. (1995). Naturalistic stimuli increase the rate and efficiency of information transmission by primary auditory afferents. Proceedings. Biol. Sci. 262, 259–265. doi:10.1098/rspb.1995.0204.

Rossant, C., Kadir, S. N., Goodman, D. F. M., Schulman, J., Hunter, M. L. D., Saleem, A. B., et al. (2016). Spike sorting for large, dense electrode arrays. Nat. Neurosci. 19, 634–641. doi:10.1038/nn.4268.

Schäfer, P. J., Corona-Strauss, F. I., Hannemann, R., Hillyard, S. A., and Strauss, D. J. (2018). Testing the limits of the stimulus reconstruction approach: auditory attention decoding in a four-speaker free field environment. Trends Hear. 22, 2331216518816600. doi:10.1177/2331216518816600.

Schwartz, O., Chichilnisky, E. J., and Simoncelli, E. P. (2002). Characterizing neural gain control using spike-triggered covariance. Adv. Neural Inf. Process. Syst., 3–5. doi:10.7551/mitpress/1120.003.0039.

Schwartz, O., Pillow, J. W., Rust, N. C., and Simoncelli, E. P. (2006). Spike-triggered neural characterization. J. Vis. 6, 484–507. doi:10.1167/6.4.13.

Sharpee, T., Rust, N. C., and Bialek, W. (2004). Analyzing neural responses to natural signals: maximally informative dimensions. Neural Comput. 16, 223–250. doi:10.1162/089976604322742010.

Stanley, G. B., Li, F. F., and Dan, Y. (1999). Reconstruction of natural scenes from ensemble responses in the lateral geniculate nucleus. J. Neurosci. 19, 8036–8042. doi:10.1523/JNEUROSCI.19-18-08036.1999.

Theunissen, F. E., Sen, K., and Doupe, A. J. (2000). Spectral-temporal receptive fields of nonlinear auditory neurons obtained Using natural sounds. J. Neurosci. 20, 2315 LP – 2331. doi:10.1523/JNEUROSCI.20-06-02315.2000.

Vahidi, N. W. (2019). Tools to investigate composite receptive fields in songbird auditory region. UC San Diego Electron. Theses Diss. Available at: https://escholarship.org/uc/item/2n08c9x0.

Vahidi, N. W., Rudraraju, S., Castagnola, E., Cea, C., Nimbalkar, S., Hanna, R., et al. (2020). Epi-Intra neural probes with glassy carbon microelectrodes help elucidate neural coding and stimulus encoding in 3D volume of tissue. J. Neural Eng. 17, 46005. doi:10.1088/1741-2552/ab9b5c.

van der Maaten, L., and Hinton, G. (2008). Visualizing data using t-SNE. J. Mach. Learn. Res. 9, 2579–2605. doi:10.1007/s10479-011-0841-3.

Velliste, M., Perel, S., Spalding, M. C., Whitford, A. S., and Schwartz, A. B. (2008). Cortical control of a prosthetic arm for self-feeding. Nature 453, 1098–1101. doi:10.1038/nature06996.

Wu, S., Amari, S.-I., and Nakahara, H. (2002). Population coding and decoding in a neural field: a computational study. Neural Comput. 14, 999–1026. doi:10.1162/089976602753633367.

Yildiz, I. B., Mesgarani, N., and Deneve, S. (2016). Predictive ensemble decoding of acoustical features explains context-dependent receptive fields. J. Neurosci. 36, 12338–12350. doi:10.1523/JNEUROSCI.4648-15.2016.

Zion Golumbic, E. M., Ding, N., Bickel, S., Lakatos, P., Schevon, C. A., McKhann, G. M., et al. (2013). Mechanisms underlying selective neuronal tracking of attended speech at a “cocktail party.” Neuron 77, 980–991. doi:10.1016/j.neuron.2012.12.037.

