## Supplemental Materials for "Spatial and Temporal Organization of Composite Receptive Fields in the Songbird Auditory Forebrain"

### *Supplementary Material*

#### 1 Supplementary Tables

| Subjects | Recording location | Cells |
|---|---|---|
| 1 | Pen 1_Lft_AP2000_ML700_Z1900<br>Pen 2_Lft_AP2000_ML700_Z2200 | 30 (4S &26M)<br>10 (3S &17M) |
| 2 | Pen 1_Lft_AP20_ML1700_Z2500<br>Pen 2_Lft_AP20_ML1680_Z2581 | 10 (3S &7M)<br>30 (9S & 21M) |
| 3 | Pen 1_Lft_AP2500_ML500_Z3200 | 8 (5S & 3M) |
| 4 | Pen 1_Lft_AP750_ML1750_Z1100<br>Pen 2_Lft_AP750_ML1750_Z1500<br>Pen 3_Lft_AP750_ML1750_Z2000 | 10 (6S & 4M)<br>30 (2S & 28M)<br>10 (3S &7M) |
| 5 | Pen 1_Lft_AP500_ML500_Z1700 | 6 (4S & 2M) |

**Table S1: Subjects and recordings used in this study.** First column: Identity of each individual starling. Second column: Number of electrode penetrations (Pen) per subject and the corresponding recording locations. Third column: Numbers of cells tested with each penetration, either single cells (S) or multi-units (M).

#### 2 Supplementary Figures

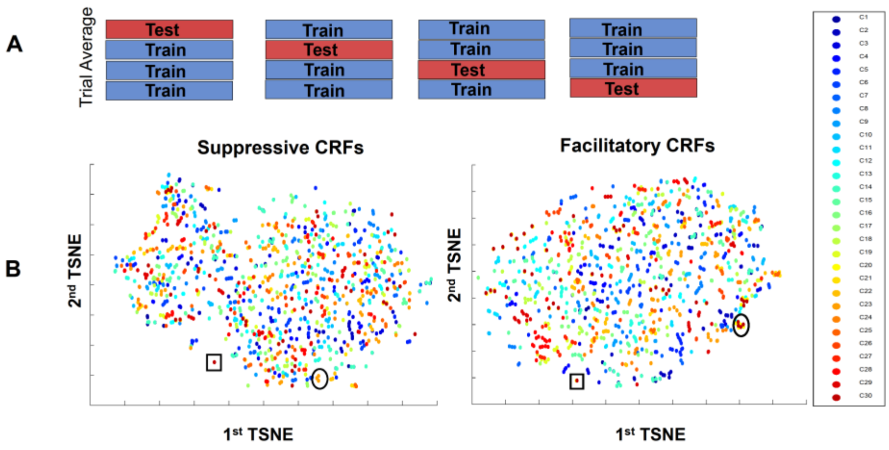


**Figure S1: Identifying noisy trials using t-SNE.** (A) Demonstration of our resampling method across the trial averages. (B) Right: Clustering similar CRFs using t-SNE for 30 cells taken from Subject 2, Penetration 2. There were 10 facilitatory features across the four resampled data sets. As an example, a set of four related CRFs are indicated with a black circle. The noisy CRFs appear as outliers from this cluster and mostly appear as a single dot. One such outlier is framed by the black square. Left: The same clustering technique repeated for suppressive CRFs. Each color indicates a different cell, and cells are ordered based on their depth along the dorsoventral plane in the NCM.

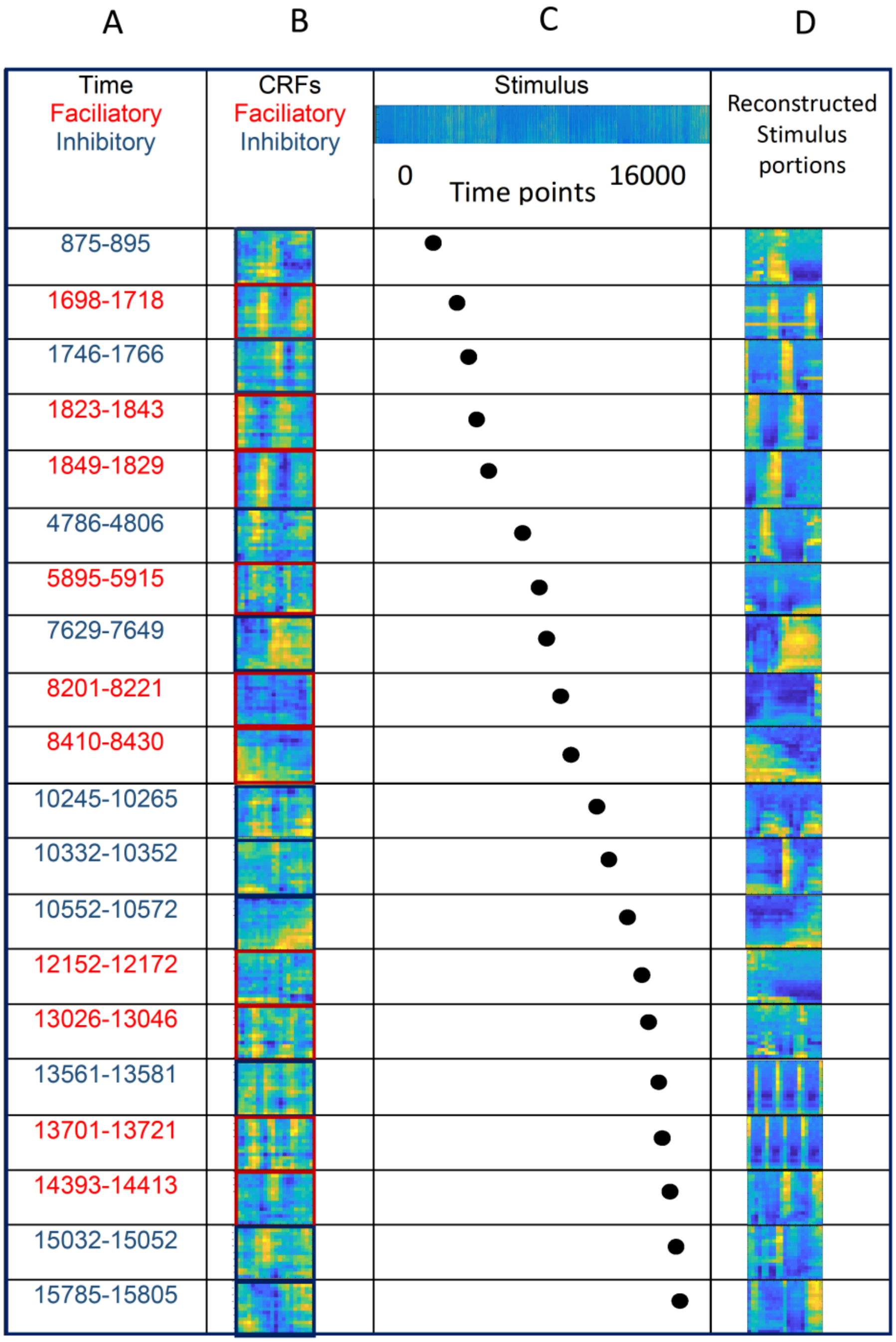


**Figure S2: Temporal map of CRFs with respect to stimuli.** (A) The 20 binned time windows where each CRF exhibited peak activation relative to a portion of the stimuli. Time windows in red are related to facilitatory responses, and time windows in blue are related to suppressive responses. (B) 20 CRFs extracted from one cell (Cell #1 from the second penetration in Subject 2), showing 10 facilitatory and 10 suppressive responses. (C) Top: Spectrogram of three concatenated bird songs. Bottom: Temporal locations of the CRFs relative to their peak activation with the stimuli (r >60%). (D) The portions of the stimuli that correspond to the CRFs, based on the temporal locations of the CRFs.
